# Slow Calcium Removal Prolongs Ventricular Relaxation in Mice with HFpEF

**DOI:** 10.1101/2025.09.10.675427

**Authors:** Tamzin Zawadzki, Diana S Usai, Jens CB Jakobsen, Morten B Thomsen

## Abstract

**Background:** Heart failure with preserved ejection fraction (HFpEF) accounts for over half of heart failure cases, yet its underlying mechanisms incompletely understood and effective therapies are lacking. Diastolic dysfunction, the hallmark of HFpEF, may arise from impaired active myocardial relaxation, but the contribution of intracellular calcium (Ca^2+^) handling remains unclear.

**Methods:** We used a validated “two-hit” murine model of HFpEF, induced by diet-driven obesity and hypertension, to investigate ventricular Ca^2+^ dynamics. Cardiac function was assessed *in vivo* by echocardiography, *ex vivo* in isolated working hearts, and at the cellular level using Fura-2-based Ca^2+^ imaging of isolated ventricular myocytes.

**Results:** HFpEF mice developed obesity, diastolic dysfunction, hypertrophy, reduced cardiac index, and exercise intolerance despite preserved ejection fraction. Impaired lusitropy was evident *in vivo, ex vivo*, and at the cellular level, where ventricular myocytes from HFpEF hearts displayed elevated diastolic [Ca^2+^]_i_, increased Ca^2+^ transient amplitudes, and frequency-dependent slowing of Ca^2+^ clearance (τ), most pronounced at 4 Hz (33% slower vs. controls, p < 0.05). HFpEF myocytes also exhibited an attenuated *β*-adrenergic response to isoprenaline, further limiting diastolic reserve.

**Conclusions:** HFpEF is characterised by a distinct ventricular myocyte Ca^2+^ handling phenotype, diverging from heart failure with reduced ejection fraction (HFrEF), with elevated diastolic Ca^2+^, exaggerated and prolonged Ca^2+^ transients, and blunted β-adrenergic modulation. These abnormalities converge to impair lusitropy and exercise tolerance, highlighting altered Ca^2+^ dynamics as a central mechanism in HFpEF. Targeting these specific Ca^2+^ handling defects may represent a novel therapeutic strategy.

## Introduction

Heart failure with preserved ejection fraction (HFpEF) has now surpassed heart failure with reduced ejection fraction (HFrEF) as the dominant form of heart failure worldwide^1,2^ creating an urgent need for mechanistically targeted therapies. Current treatments, largely developed for HFrEF, fail in HFpEF, underscoring its distinct pathophysiology^3,4^. This problem is compounded by the profound heterogeneity of HFpEF; patients fall into diverse ‘phenogroups’ characterized by various combinations of comorbidities such as obesity, diabetes, hypertension and systemic inflammation^1,4,5^ – all of which can independently impair cardiac function. Such heterogeneity is impossible to recapitulate in a single preclinical model. Existing approaches either reproduce isolated features or rely heavily on inducing comorbidities^6–13^ confounding primary myocardial abnormalities with systemic effects. Consequently, the mechanisms driving diastolic dysfunction, the diagnostic hallmark of HFpEF, remain incompletely understood.

Insights into HFpEF are additionally limited by the scarcity of myocardial biopsies, as these patients undergo cardiac surgery far less frequently than those with HFrEF. Diastolic dysfunction has traditionally been attributed to reduced myocardial compliance caused by extracellular matrix (ECM) stiffening due to excess collagen deposition^14^. However, the largest biopsy study of HFpEF found significant fibrosis in only 27% of patients^15^, indicating that in most cases non-fibrotic mechanisms impair diastole. Comparing two cohorts of HFpEF patients, Rosch *et al*. demonstrated that those with left ventricular ejection fraction (LVEF) >60% exhibit higher diastolic stiffness despite lower extracellular volume fractions than those with 50-60% LVEF, suggesting stiffness may arise from intracellular mechanisms^15^. Evidence from isolated cardiomyocytes supports this view, suggesting an intracellular origin of dysfunction. For example, in chemically skinned cardiomyocytes from patients with HFpEF, Borbély *et al*. demonstrated that the demembranated cells are inherently stiff. Elevated passive tension was normalized following PKA administration, implicating hypophosphorylation of sarcomeric proteins as a causal factor in stiffness^16^. Multiple studies have confirmed post-translational modifications to sarcomeric proteins, particularly titin, as important determinants of cardiomyocyte stiffness^9,17–20^.

However, diastolic relaxation (lusitropy) *in vivo* is not solely a passive process, it is a dynamic event governed by both active and passive components. Calcium-dependent lusitropy refers to the active, energy-dependent process of myocardial relaxation, whereby the removal of cytosolic calcium during diastole promotes cross-bridge detachment and facilitates ventricular relaxation^21^. Selby *et al*. showed that in myocardial strips isolated from HFpEF patients, reduced sodium–calcium exchanger (NCX) reserve limits calcium extrusion. During tachycardia this overwhelms the sarcoplasmic reticulum (SR)’s calcium sequestration capacity, hampering lusitropy and resulting in incomplete relaxation due to calcium overload^22^. Ostensibly diminished calcium-dependent lusitropy could exacerbate passive stiffness from pathologically altered myofilaments further compromising diastolic function.

We hypothesize that pathological calcium handling is a key cellular origin of diastolic dysfunction in HFpEF. Using a mouse model that recapitulates the clinical and hemodynamic features of heart failure, but with preserved ejection fraction, we identify a distinct pattern of frequency-dependent calcium dysregulation in ventricular myocytes. We establish a mechanistic continuum from cellular dysregulation to whole-organ dysfunction, showing that these calcium handling abnormalities manifest as impaired lusitropy in both *ex-vivo* working hearts and *in vivo*. Furthermore, we find that HFpEF myocytes exhibit a blunted *β-*adrenergic response, directly implicating this specific defect in the reduced diastolic reserve and exercise intolerance that is characteristic of both our HFpEF animal model and the clinical syndrome. By identifying impaired calcium handling as a driver of pathophysiology, our work highlights impaired lusitropy as a therapeutic target in a disease urgently requiring effective treatment.

## Results

### Mice on HFpEF protocol develop progressive obesity

Over the 16-week protocol (Fig. 1A), body mass was recorded every two weeks, shown in Fig. 1B. All mice gained body mass commensurate with age (mixed-effects model, p<0.0001 for time; Fig. 1B), with HFpEF mice exhibiting a higher mean body mass compared with the controls (Fig. 1B). Consequently, HFpEF mice had a greater total body mass gain than controls (17.6 ± 5.4 g vs. 7.4 ± 3.1 g; p < 0.0001, unpaired t-test; Fig. 1C), which over the duration of the protocol corresponded to a 77 ± 21% increase in body mass for HFpEF mice compared with only 31 ± 13% in controls.

**Figure 1.**
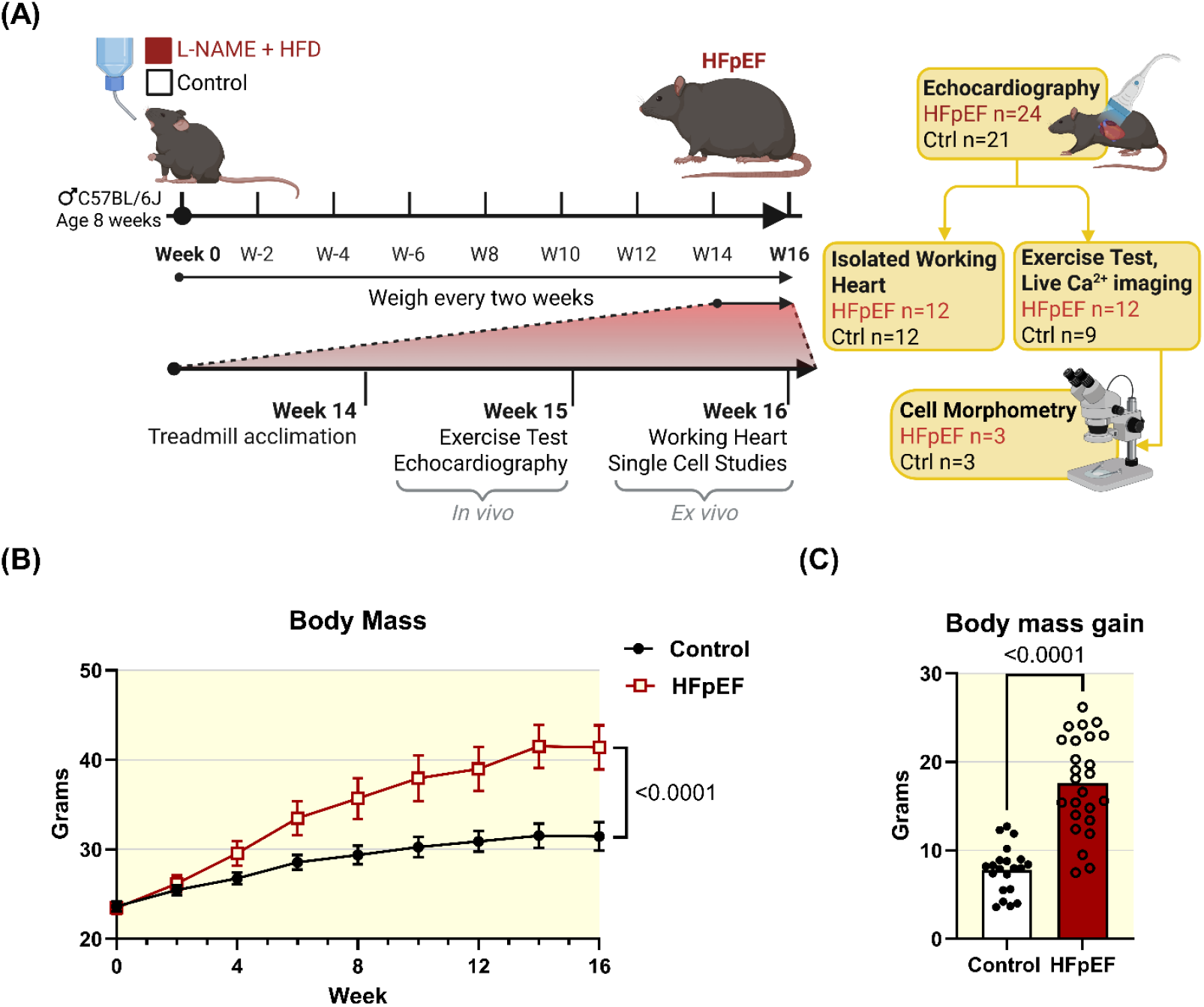
HFpEF mice develop higher body mass than controls over 16 weeks. (A) Schematic of the experimental timeline showing treatment conditions, key timepoints, and group sizes for each protocol. (B) Body mass progression over 16 weeks is consistently higher in the HFpEF (n=24) group compared with controls (n=21). Mean ± 95% CI. Statistical analysis: mixed-effects model with time as a repeated measure; *p* value shown in the figure refers to the main effect of group (HFpEF vs. control). (C) Total body mass gain from week 0 to 16 was significantly higher in HFpEF mice than in controls. Statistical analysis: unpaired Student’s t-test, *p* value shown in figure.

### HFpEF mice develop diastolic dysfunction *in vivo* with preserved ejection fraction

Cardiac function was evaluated by transthoracic echocardiography under anaesthesia after 16 weeks on the HFpEF protocol (Fig. 2). Ejection fraction in the HFpEF mice was preserved, showing no significant difference from control animals (Fig 2A). To evaluate cardiac performance relative to body size, we determined cardiac index (CI) – cardiac output normalized to body surface area. Given the substantial increase in body mass induced by the high-fat diet (Fig 1), CI provided a more physiologically relevant metric of cardiac function than cardiac output. The HFpEF group had a significant reduction in CI compared to the control group (Fig 2B), indicating progression to HF despite preserved EF and no change in heart rate (Fig 2C). Diastolic function was assessed by measuring E/e’ and the isovolumetric relaxation time (IVRT). E/e’, which is directly linked to LV filling pressure, was significantly increased in the HFpEF group compared with controls (Fig 2D). The duration of the isovolumic relaxation of the LV was prolonged by 17.4 % in the HFpEF group compared with controls (Fig. 2E). The LV wall was measured in diastole and showed a 13.4 % increase in thickness in the HFpEF group compared to controls, indicating left ventricular hypertrophy (Fig 2F). Collectively, the echocardiographic data show that mice subjected to obesity and hypertension for 16-weeks develop a phenotype consistent with HFpEF, characterized by preserved ejection fraction, diastolic dysfunction, and left ventricular hypertrophy.

**Figure 2.**
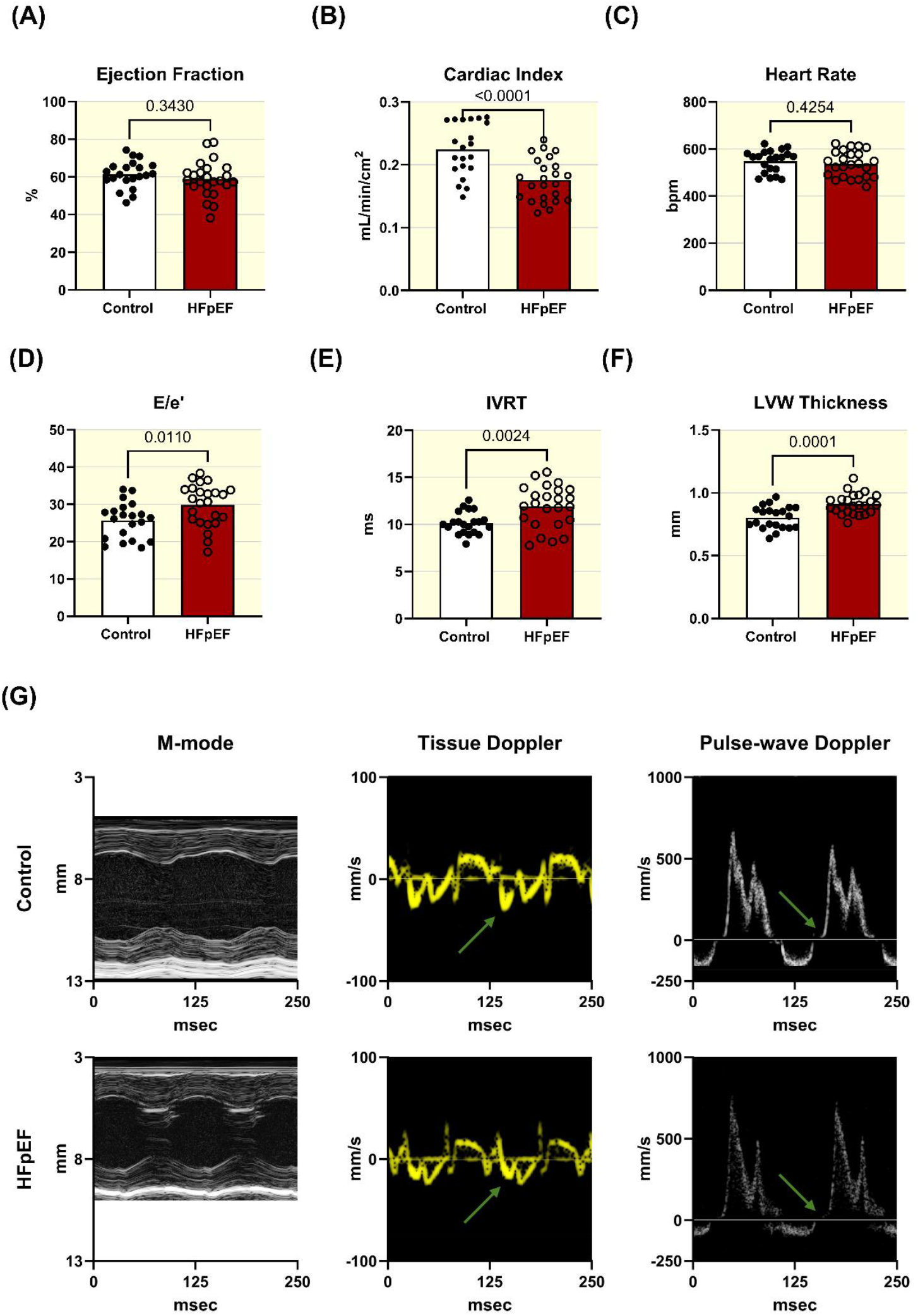
HFpEF mice exhibit diastolic dysfunction *in vivo*, as measured by echocardiography. Echocardiographic assessment after the 16-week HFpEF protocol in HFpEF (n = 24) and control (n = 21) mice. Graphs show individual data points per mouse. Statistical analyses: unpaired Student’s t-tests, *p* values shown in figure panels. (A) Ejection fraction was preserved in HFpEF mice. (B) Cardiac index (cardiac output relative to body-surface area) was significantly reduced in the HFpEF compared with control group. (C) Heart rate during the echocardiographic assessment was similar between groups. (D) E/e′ ratio was increased in HFpEF mice compared to controls. (E) Isovolumic relaxation time (IVRT) was prolonged in the HFpEF group. (F) Left ventricular wall (LVW) thickness was significantly increased in HFpEF mice. (G) Representative echocardiographic images from a control (top) and HFpEF (bottom) mouse. Left: M-mode images showing the anterior and posterior free wall of the LV. Middle: tissue Doppler signals showing tissue velocity. The green arrows show the e’ wave during LV filling. Right: Pulse-wave Doppler trace showing blood velocity at the level of the mitral valve. The green arrows point to the isovolumic relaxation phase that is prolonged in HFpEF.

### Clinical hallmark of exercise intolerance recapitulated in HFpEF mice

Exercise intolerance is a clinical feature of heart failure and is essential for establishing translational relevance in preclinical models. To assess this, mice underwent exercise testing (Fig. 3A). By week 15, HFpEF mice exhibited marked exercise intolerance, running a mean distance of 175 ± 8 meters until exhaustion—a 37% reduction compared to control mice, which ran 257 ± 23 meters (Fig. 3B).

**Figure 3.**
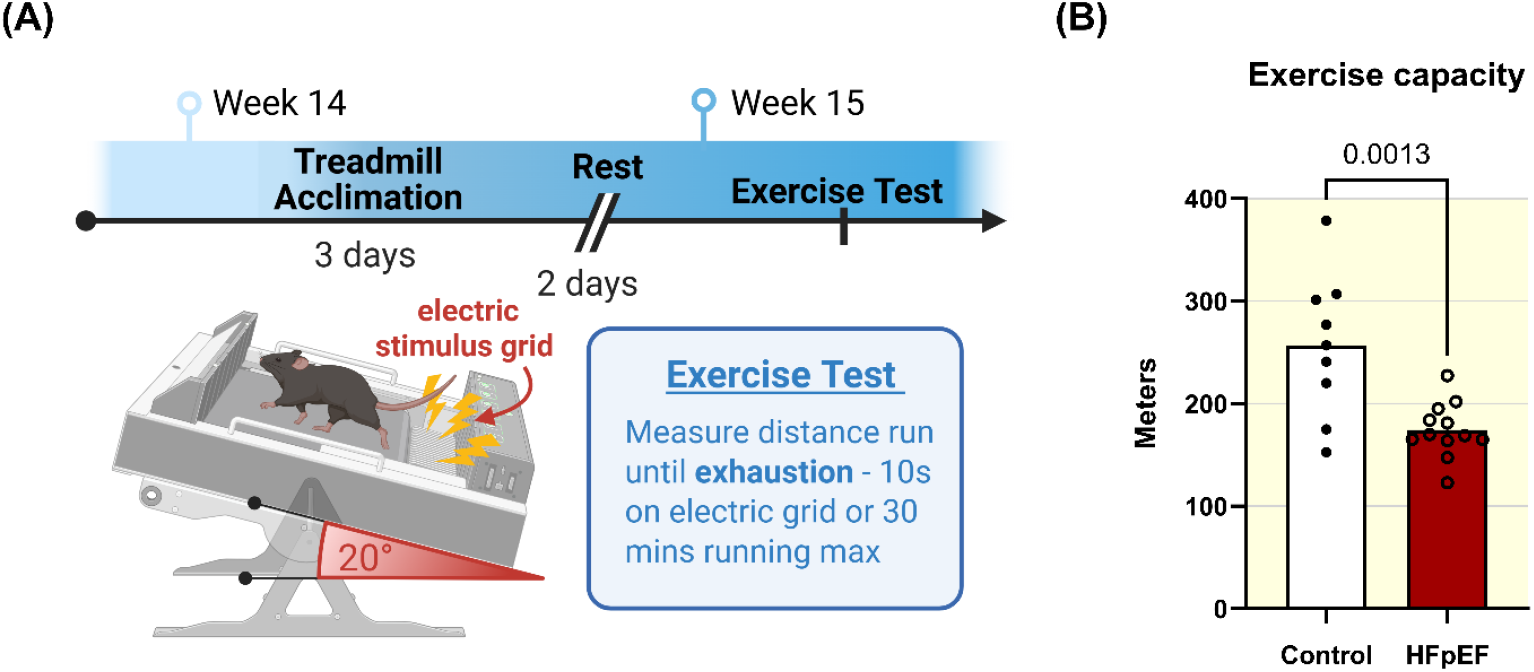
HFpEF mice exhibit exercise intolerance. (A) Schematic illustrating the protocol used to assess exercise intolerance at week 15. Acclimation to the treadmill for 3 consecutive days, followed by a test for exercise capacity after 2 additional ‘rest’ days. (B) HFpEF mice (n = 12) ran a shorter mean distance to exhaustion compared to controls (n = 9). Statistical analysis: Student’s unpaired t-test, *p* value shown in figure.

### Diastolic Dysfunction *ex vivo* in isolated HFpEF hearts

Cardiac function was assessed using an isolated working heart setup, enabling evaluation of myocardial performance under controlled preload conditions without influence from *in vivo* compensatory mechanisms^23^. Hearts from HFpEF animals exhibited significantly reduced cardiac output compared to controls (Fig. 4A). These recordings were made under constant heart rate, so stroke volume was also significantly lower in hearts from HFpEF mice (data not shown). No difference in peak LV pressure was observed between groups (Fig. 4B). The peak negative slope of the ventricular pressure curve during relaxation (–dP/dt) was significantly reduced in HFpEF hearts, indicating slower and impaired ventricular relaxation (Fig. 4C). This was consistent with a significantly prolonged LV diastolic time constant, τ, in HFpEF hearts compared with controls (Fig. 4D). Together, these findings corroborate impaired diastolic relaxation in HFpEF hearts. Moreover, these *ex-vivo* results reinforce the *in vivo* echocardiographic evidence of an impairment of the ventricular mechanical relaxation observed in HFpEF mice (Fig 2D and E).

**Figure 4.**
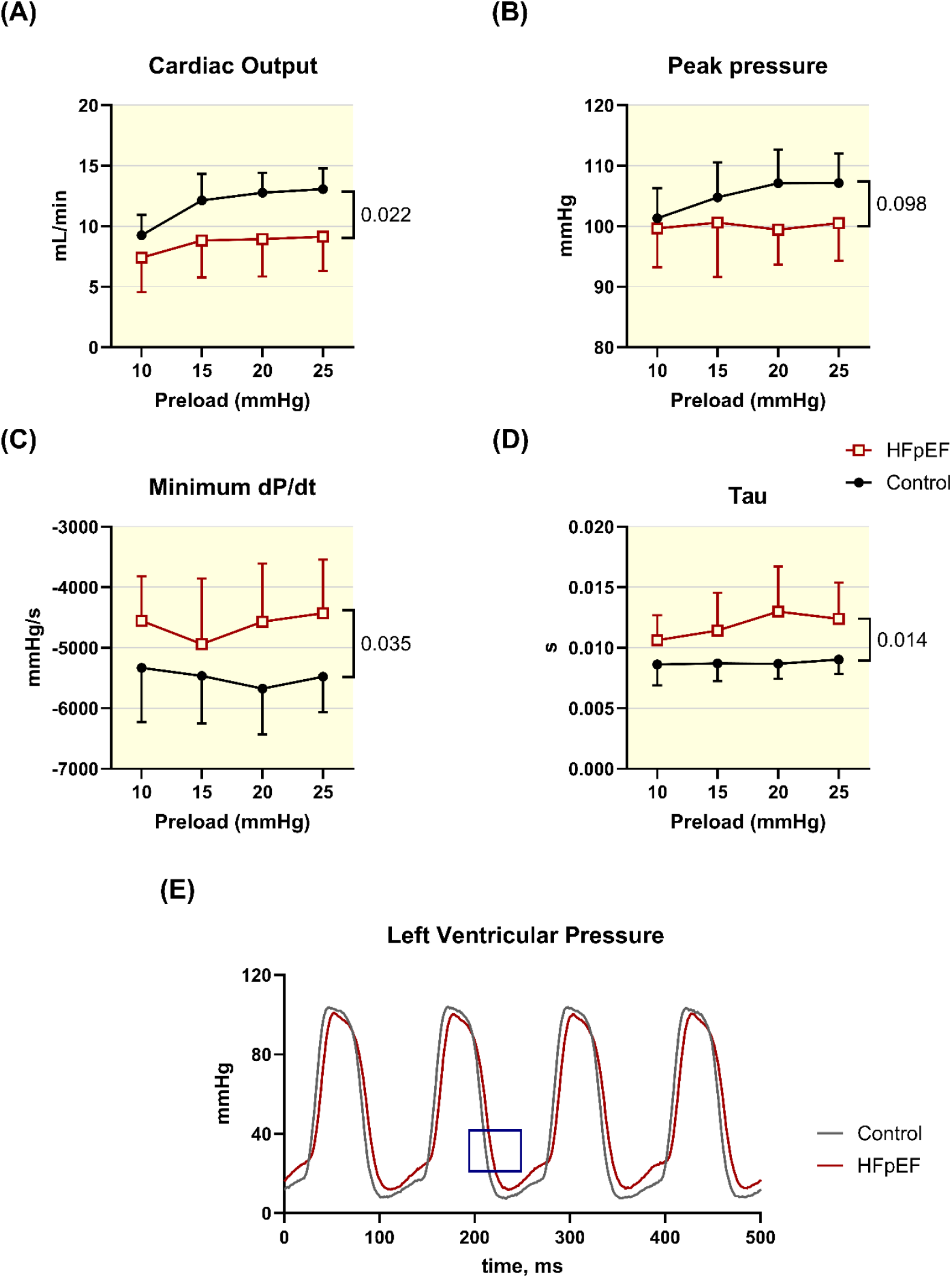
HFpEF mice exhibit diastolic dysfunction *ex vivo*, as measured in isolated working hearts. Hearts isolated from HFpEF mice (n= 9) and from age-matched control mice (n = 8) at week 16. Hemodynamic responses reported across a range of filling pressures (preload: 10–25 mmHg).Statistical analyses: two-way ANOVA, with factors for group (HFpEF vs. control) and preload. *P* values reported in the figure refer to the main effect of group. (A) Cardiac output was significantly lower in hearts from HFpEF mice compared to controls. Heart rate was kept constant at 480 beats per minute. (B) Peak LV pressure during sinus rhythm similar in HFpEF and control hearts. (C) The peak negative slope of the pressure curve during ventricular relaxation (–dP/dt) was less steep in HFpEF than in control hearts. (D) A mono-exponential fit to ventricular pressure in the relaxation phase (from peak –dP/dt to end-diastolic pressure) was used to derive the relaxation time constant, τ. This was significantly higher in HFpEF compared with control hearts. (E) Representative 0.5 sec trace of the LV pressure at 20 mmHg preload in a HFpEF (red line) and a control (grey line) heart. Note the slower ventricular relaxation. The blue rectangle indicates the relaxation phase used to calculate τ (panel D).

### Cellular Hypertrophy Underlies Echocardiographic LV Wall Thickening in HFpEF mice

Consistent with increased LV wall thickness observed in the hearts of HFpEF mice (Fig 2F), ventricular myocytes isolated from HFpEF hearts exhibited significant hypertrophy at the cellular level. Left ventricular myocytes from HFpEF hearts were 12% longer, 17% wider, and displayed 25% larger cross-sectional area compared to controls (Fig. 5).

**Figure 5.**
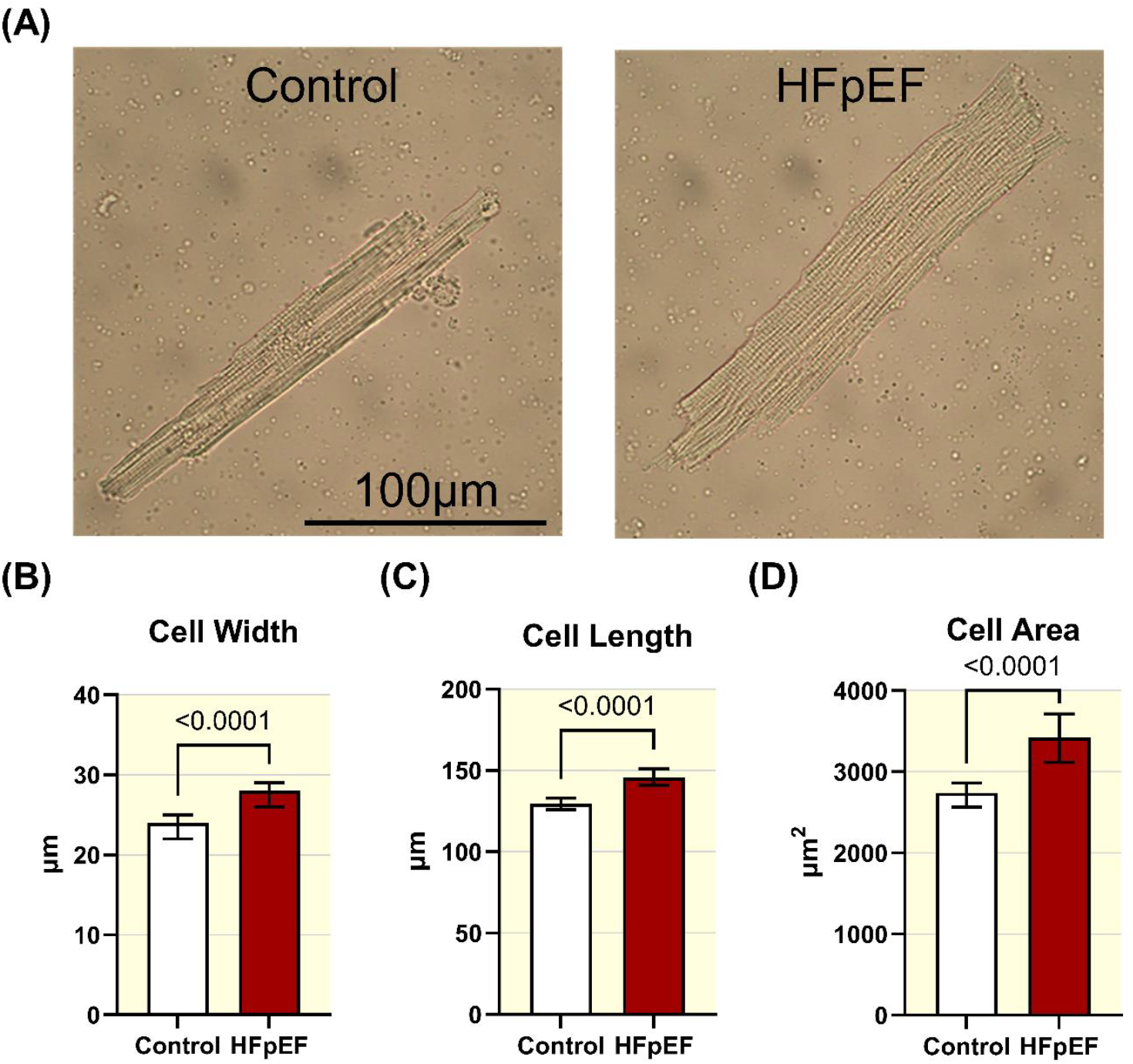
Ventricular myocytes from HFpEF hearts are hypertrophic. (A) Representative images of isolated myocytes from control (left) and HFpEF (right) mice. (B–D) Quantification of cell width, length and cross-sectional area, showing significantly increased dimensions in HFpEF myocytes (n = 134 cells, 3 hearts) compared to controls (n = 189 cells, 3 hearts). (B) Cell width (mean ± 95% CI; normally distributed according to Shapiro-Wilk test; *p*-value from unpaired Student’s t-test reported in figure). (C) Cell length (median ± 95% CI; not normally distributed, Shapiro-Wilk test; *p*-value from Mann-Whitney reported in figure). (D) Cross-sectional area (median ± 95% CI; not normally distributed, Shapiro-Wilk test; *p*-value from Mann-Whitney reported in figure).

### Ventricular myocytes from HFpEF hearts display impaired calcium handling with frequency-dependent abnormalities

*In vivo* echocardiographic assessment revealed prolonged LV relaxation in HFpEF hearts (Fig. 2E), indicative of diastolic dysfunction. This finding was corroborated *ex vivo* in isolated working heart analyses, where hearts from HFpEF mice demonstrated a significantly prolonged LV diastolic time constant (τ; Fig. 4D) and reduced peak negative dP/dt (Fig. 4C), even under tightly controlled LV filling pressure. Impaired relaxation in the absence of systemic influences strongly suggests an intrinsic myocardial defect in HFpEF hearts. Given diastolic relaxation is contingent upon efficient cytosolic Ca^2+^ clearance, we next investigated whether aberrant Ca^2+^ handling contributes to the observed dysfunction.

Ventricular myocytes from HFpEF hearts displayed dysregulated Ca^2+^ handling compared to controls, shown in Fig. 6. Fig. 6A shows representative Ca^2+^ transients from cells from HFpEF and control hearts at two different pacing frequencies. Diastolic Ca^2+^_i_(R_0_) was tonically elevated by 7% in HFpEF myocytes at 0.5 Hz, with a tendency to remain elevated at higher stimulation frequencies but without reaching statistical significance (Fig. 6B). Median CaT amplitude (R/R_0_) was significantly elevated in HFpEF at all stimulation frequencies except at 1 Hz (Fig. 6B). Time to peak was significantly prolonged in HFpEF compared with control cells at stimulation frequencies of 0.5–3 Hz (Fig. 6B). Median τ tended to be higher in HFpEF than control cells, reaching statistical significance at 4Hz stimulation, where median τ was 33% higher in HFpEF compared with controls. Higher τ indicates slower Ca^2+^ removal from the cytosol. A non-linear regression analysis was performed to compare τ in control and HFpEF cells across stimulation frequencies, shown Fig 6.C. The regression curves showed that τ decreases in both groups at higher stimulation frequencies, indicating a frequency-dependent acceleration in Ca^2+^ removal. Interestingly, comparison of the two curves using a sum-of-squares F-test revealed a significant difference, indicating a frequency-dependent impairment in Ca^2+^ removal in HFpEF relative to control cells (Fig. 6C).

**Figure 6.**
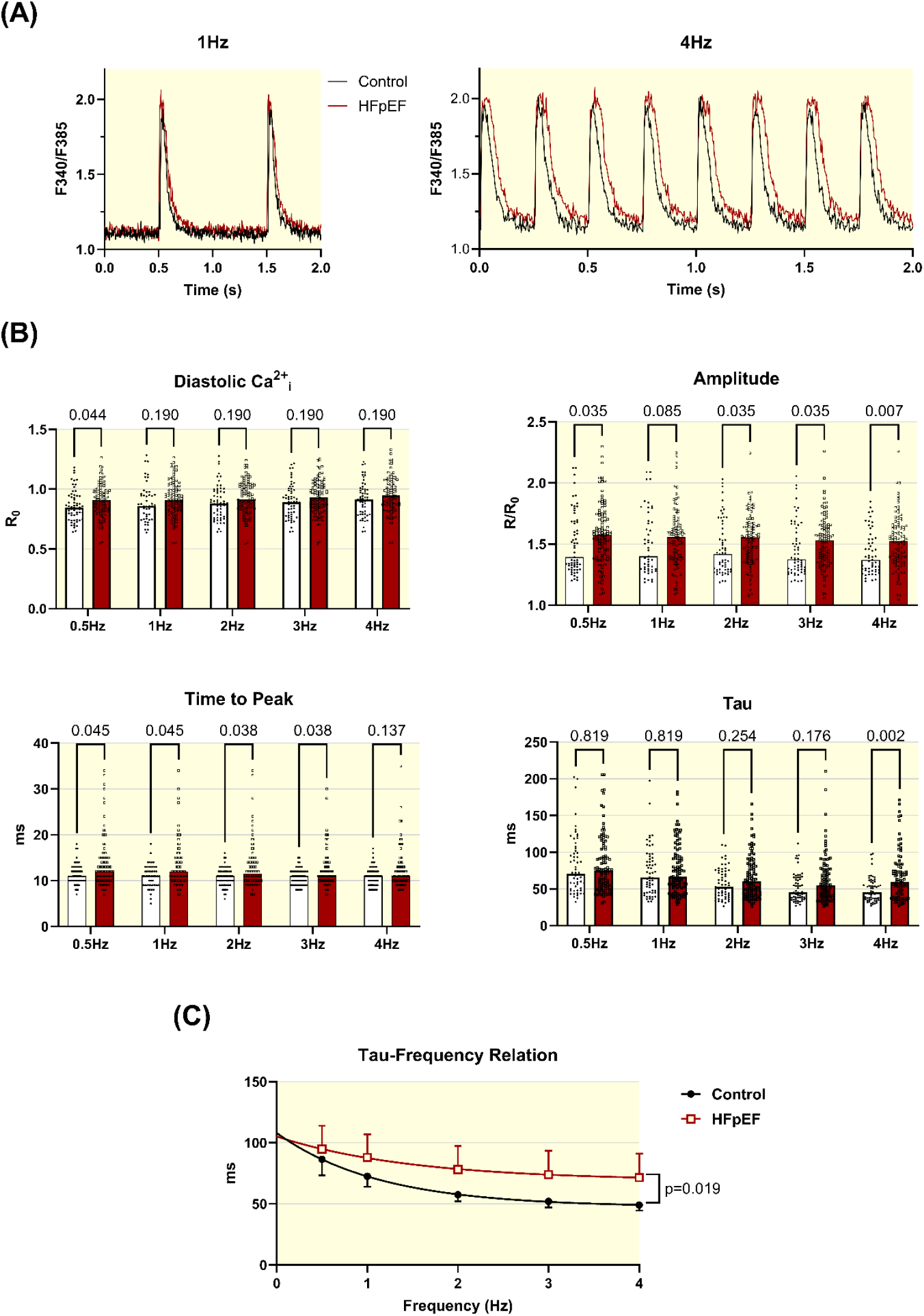
Ventricular myocytes from HFpEF hearts exhibit frequency-dependent calcium handling dysfunction. (A) Representative Fura-2 fluorescence traces showing the ratiometric (R) signal (F_340_/F_385_), which reflects free intracellular calcium concentration normalized fordye loading and cell thickness. Traces are shown for stimulation at 1 Hz (left) and 4 Hz (right). (B) Quantification of calcium transient (CaT) parameters: diastolic Ca^2+^_i_ (baseline| or R_0_), amplitude (R/R_0_), time-to-peak, and tau (τ) across 0.5-4 Hz stimulation frequencies. Data from control (n = 61 cells, 4 hearts) and HFpEF (n = 106 cells, 8 hearts) myocytes. Compared with controls, HFpEF myocytes showed: significantly elevated diastolic Ca^2+^_i_ (at 0.5 Hz only), higher CaT amplitude (all frequencies except 1 Hz), prolonged time to peak (0.5-3 Hz), and prolonged τ at 4 Hz. Statistics: Data were non-normally distributed (Shapiro–Wilk test); median values were compared using Mann– Whitney tests with Holm–Šídák correction for multiple comparisons. Adjusted *p*-values are reported in the figure. (C) τ across stimulation frequencies were fitted with a one-phase decay model, demonstrating frequency-dependent acceleration of Ca^2+^ clearance, which was impaired in HFpEF myocytes (sum-of-squares F-test). Data are mean ± 95% CI. P value in graph shows that the two fitted curves are statistically different.

### Attenuated isoprenaline response in HFpEF cardiomyocytes

Given the frequency-dependent impairment in Ca^2+^ reuptake kinetics observed in HFpEF myocytes (Fig. 6) and the exercise intolerance exhibited by HFpEF mice (Fig. 3), we sought to determine whether these defects were associated with a blunted *β*-adrenergic response. To address this, we examined the effect of an inotropic intervention using isoprenaline. Fig. 7A shows representative Ca^2+^ traces before and after isoprenaline in a HFpEF and a control myocyte. Ca^2+^ transient (CaT) decay time, defined as the time to reach 90% decay from the peak amplitude, was measured in the same cells before and after addition of isoprenaline to the superfusion solution. Isoprenaline reduced decay time in both groups (paired measurements; Fig. 7B), but the reduction was significantly smaller in HFpEF compared with control myocytes at every stimulation frequency. The difference was most pronounced at 0.5 Hz, where isoprenaline accelerated τ by 33% in control myocytes but only by 18% in HFpEF myocytes (Fig. 7C). This difference persisted across all stimulation frequencies, being more pronounced at 0.5–2 Hz stimulation (p ≤ 0.05) and less so at 3 and 4 Hz (p ≥0.05). Together,these findings indicate a mechanistic defect in *β*-adrenergic modulation of Ca^2+^ reuptake in HFpEF myocytes, which may contribute to impaired diastolic reserve and exercise intolerance observed *in vivo*.

**Figure 7.**
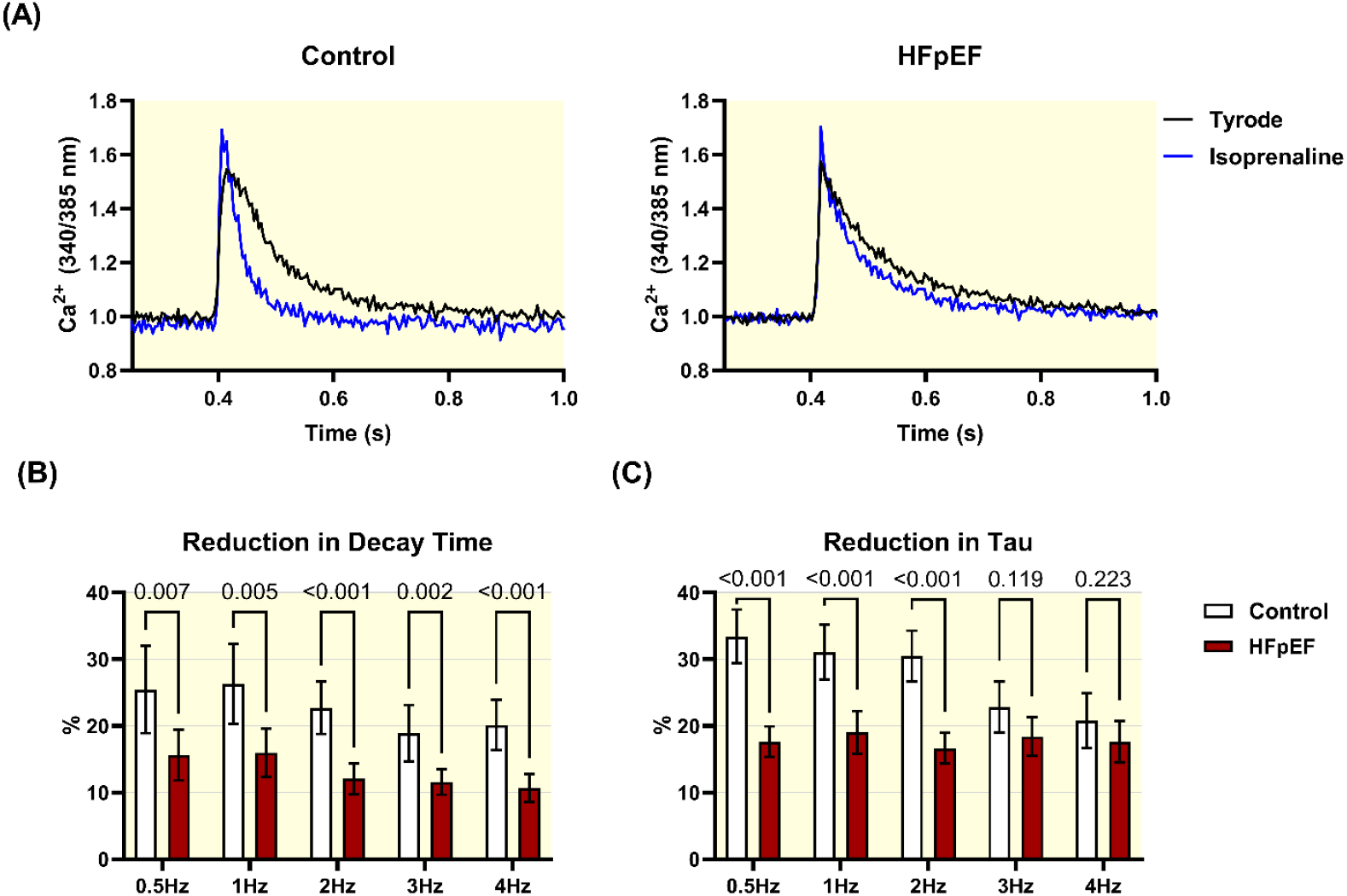
HFpEF ventricular myocytes show impaired isoprenaline response. (A) Representative Fura-2 traces from control (left) and HFpEF (right) myocytes stimulated at 0.5 Hz in Tyrode solution (black) and after 2 minutes in 50 nM isoprenaline (blue). (B) Paired data showing the isoprenaline-induced reduction in calcium transient decay time to 90% decay. HFpEF myocytes exhibit a significantly smaller reduction in decay time following isoprenaline treatment compared to controls. (C) Paired data showing the isoproterenol-induced reduction in τ. HFpEF myocytes exhibit a significantly smaller reduction in τ following isoprenaline treatment compared to controls at stimulation frequencies of 0.5–2 Hz. Sample sizes: HFpEF myocytes (n = 48 cells, 4 hearts) and control (n = 36 cells, 4 hearts). Data are median ± 95% CI. Distributions were non-normal by Shapiro–Wilk test, statistical comparisons used Mann–Whitney tests corrected for multiple comparisons (Holm–Šídák method). Adjusted *p*-values are reported in the figure.

## Discussion

Abnormalities in calcium signalling are a unifying mechanism in the pathogenesis of HF^24–26^. Whilst Ca^2+^ dyshomeostasis is well-studied in the context of HFrEF^21,26–33^, less is known regarding its role in the aetiology of HFpEF^34^. Fibrosis, metabolic stress, and systemic inflammation have dominated HFpEF research^9,17,35–39,^ whereas calcium handling is comparatively neglected^12,40,41^. This gap is striking, given the fundamental dependence of diastole on spatiotemporally co-ordinated calcium cycling^21,42^. Pathophysiological calcium handling may therefore represent a common myocardial mechanism that spans the otherwise diverse HFpEF phenogroups, offering a viable therapeutic target.

Here we describe a pattern of frequency-dependent Ca^2+^ dysregulation in ventricular myocytes from a validated “two-hit” murine model of HFpEF^11,43^. Mice subjected to diet-induced obesity and hypertension maintained a normal ejection fraction but exhibited diastolic dysfunction, hypertrophy, reduced cardiac index, and exercise intolerance - recapitulating the clinical syndrome. Critically, diastolic dysfunction, the defining feature of HFpEF, was reproduced from whole animal to single myocytes: prolonged myocardial relaxation and increased filling pressures *in vivo* (Fig. 2) were mirrored *ex vivo* (Fig.4), suggesting an intracardiac origin and reflecting slow diastolic calcium removal in isolated ventricular myocytes (Fig 6). Hypertrophic remodelling in HFpEF hearts detected by echocardiography (Fig. 2) was also evident in isolated myocytes (Fig. 5), implicating a cellular – rather than an extracellular or fibrotic – origin of remodelling. Concordance across our findings suggests impaired myocyte calcium handling contributes directly to diastolic dysfunction *in vivo*.

Our results align with emerging evidence that calcium dyshomeostasis impairs lusitropy, contributing to exercise intolerance in HFpEF. Slow cytoplasmic Ca^2+^ clearance delays its dissociation from myofilaments prolonging relaxation^42^. Selby *et al*. reported incomplete relaxation in LV strips from HFpEF patients at pacing rates ≥2Hz^22^, comparable to the frequency-dependent impairment in Ca^2+^ removal we observed. HFpEF myocytes showed a consistent trend toward elevated diastolic Ca^2+^ across all pacing frequencies, reaching significance at 0.5 Hz (Fig. 6). Similar findings in human HFpEF and rodent models support calcium overload as a key contributor to impaired lusitropy during tachycardia^22,40,41,44^. The most severe effect on CaT kinetics was observed at 4Hz, where Ca^2+^ clearance (τ) was 33% slower in HFpEF myocytes than in controls (Fig. 6). In a hypertensive rat model of HFpEF, Kilfoil *et al*. reported similar CaT decay times between control and HFpEF myocytes at 1 Hz^41^, but their absolute decay times were markedly slower, likely due to experiments conducted at room temperature rather than the more physiological 37 °C used here. Notably, at their highest stimulation rate (2Hz), they similarly recorded elevated diastolic calcium in HFpEF myocytes.

Frisk *et al*. also observed slow Ca^2+^ removal at 4Hz but not 1Hz, albeit in an aetiology-dependent manner, occurring in obese-but not hypertensive-HFpEF rat myocytes^40^. Conversely, Kaplan *et al*. reported an increased Ca^2+^ removal rate in a “three-hit” HFpEF mouse model, which have high dietary fructose in addition to obesity and hypertension, however this was confined to 1 Hz pacing and lost at 4 Hz^12^. While such discrepancies may arise from model-specific pathophysiology or methodological variability in Ca^2+^ handling assessments, the collective evidence consistently highlights calcium mishandling as a central mechanism in the pathogenesis of HFpEF.

We further demonstrate that myocytes from HFpEF mice exhibit an attenuated *β*-adrenergic response, showing reduced sensitivity to isoprenaline-induced acceleration of Ca^2+^ clearance (Fig. 7). This blunted response likely compounds slow cytosolic Ca^2+^ clearance at high heart rates and contributes to exercise intolerance observed *in vivo* (Fig 3), a finding consistent with the attenuated *β*-adrenergic signalling reported in rat HFpEF^41^.

Our data reveal a key divergence from the classic heart failure paradigm: rather than the canonical decline in CaT amplitude, which governs the magnitude of contraction, thus underpinning its contractile deficit in HFrEF^26,32,33^, here we observe the opposite. HFpEF myocytes exhibited significantly higher CaT amplitudes compared with controls. This aligns with preserved systolic function *in vivo* and mirrors earlier observations in HFpEF rats^40,41^. Multiple studies document a pattern of early, compensatory alterations in calcium handling which initially conserve function but ultimately become maladaptive over time, contributing to progressive dysfunction ^27,28,45,46^.Consistent with this compensatory-to-maladaptive continuum, it is tempting to speculate that exaggerated CaT amplitudes constitute a compensatory mechanism aimed at preserving contractile performance, reflecting a need for enhanced SR Ca^2+^ release to overcome pathologically elevated myocardial stiffness - characteristic of HFpEF hearts ^9,17,18,47,48^.

Our data reveals a mechanistic paradox: although larger amplitude Ca^2+^ transients in HFpEF myocytes suggest increased SR Ca^2+^ availability, Ca^2+^ transient decay (τ) is significantly prolonged. We propose that this paradox reflects competing determinants of Ca^2+^ clearance. At low stimulation frequencies (e.g., 0.5 Hz), elevated diastolic [Ca^2+^] acts as a compensatory mechanism by enhancing SERCA2a uptake, masking a reduction in SERCA Vmax and thereby minimizing changes in τ. However, at high stimulation rates, this compensatory mechanism is insufficient, unmasking the intrinsic deficit in SERCA function and producing a progressively slower τ. Such frequency-dependent impairment of Ca^2+^ clearance may critically limit diastolic relaxation under stress conditions, providing a key cellular mechanism for the impaired lusitropy and exercise intolerance characteristic of HFpEF. Persistently elevated diastolic Ca^2+^ can activate hypertrophic pathways, including calcineurin/NFAT signaling^49^, directly linking the elevated diastolic Ca^2+^ we observed to both impaired lusitropy and the development of the hypertrophy characteristic of the HFpEF phenotype.

In summary, our findings reveal a distinct constellation of intracellular Ca^2+^ dysregulation in HFpEF, which diverges from the pathophysiological patterns observed in HFrEF. Rather than reflecting a classic heart failure phenotype, HFpEF myocytes exhibit a unique combination of elevated diastolic [Ca^2+^]_i_, increased and prolonged Ca^2+^ transients, and blunted frequency- and β-adrenergic-driven modulation. Collectively, our data extend previous observations by demonstrating that impaired Ca^2+^ clearance in HFpEF myocytes becomes most evident under high-frequency stimulation, revealing a limited lusitropic reserve that likely contributes to exercise intolerance

## Methods

The experiments in this study received approval from the national ethics committee under the Danish Ministry of Environment and Food (license #2023-15-0201-01576). They adhered to the European Parliament Directive on the Protection of Animals Used for Scientific Purposes. Experiments were conducted in full compliance with the guidelines set forth by both institutional and national animal care committees.

### Animals

Forty-eight male C57BL6/NRj mice, aged 5 weeks, were obtained from Janvier Labs and used in these experiments. Mice were housed 4 animals per cage, kept at 20–22 °C in a 12-hour light/dark cycle and fed standard chow *ad libitum*. Three mice were euthanized during the acclimatisation period due to injuries sustained from fighting. After a week acclimatisation period mice, aged 8 weeks, were randomized into ‘control’ and ‘HFpEF’ groups. Control mice were started on a low-fat diet with 10% calories from fat (D12450Ji, Research Diets) and tap water *ad libitum*. HFpEF animals were fed high-fat diet with 60% calories from fat (D12492i, Research Diets) *ad libitum* and 1.0 g/L of N_ω_-Nitro-D-arginine methyl ester (L-NAME, Sigma Aldrich) dissolved in tap water^43^. Diet and water bottles were changed weekly.

### Echocardiography

Transthoracic echocardiography was performed on anaesthetised mice. After induction, the isoflurane was kept on 1.5-2.0% in O_2_. The mice were placed in supine position on a heated platform. Body temperature was monitored and kept at physiological level (36-38°C) throughout the examination. The echocardiography was conducted with an ultrasound machine designed for rodents (Vevo 3100, VisualSonics, Canada) and a transducer designed for cardiac examinations on mice (MS550D, VisualSonics, Canada).

Parasternal short axis view in b-mode and m-mode and apical four chamber view were obtained following the guidelines from European Society of Cardiology.^50^ Quantification of the data were done offline and blinded to group (HFpEF or control) with the VevoLab software (Version 5.8.1, VisualSonics, Canada). Ejection fraction was determined according to Simpson’s principle. Cardiac index was calculated as cardiac output relative to the body surface area, which was estimated from body mass. E/e’ and isovolumic relaxation time were determined from tissue and pulse-wave Doppler traces. Left ventricular wall thickness was calculated as an average of the anterior and posterior wall in diastole.

### Exercise test

Mice were acclimatised to a treadmill (Exer-3/6, Columbus Instruments) over 3 consecutive days. An exercise capacity test was conducted after 2 days of rest following the training protocol: Mice ran on an incline of 20%, starting at 6 m/min for 4 minutes, then 12 m/min. After another 4 minutes, speed was increased incrementally by 2 m/min every 3 minutes to a maximum speed of 20 m/min. Animals were run until exhaustion or for a maximum time of 30 minutes. Criteria for exhaustion was an inability to run, defined by 10 second standstill on the electric stimulus grid. Animals were returned to the home cage immediately after the point of exhaustion was reached. Total covered distance (in metres) was the outcome of the exercise capacity test.

### Isolated working heart perfusion

The isolated working-heart technique has been described in detail by us previously.^23^ Mice were euthanized by cervical dislocation, and the heart was quickly extracted into a dish. The aorta was cannulated (cannula: length = 4.1 mm; inner diameter = 0.80 mm) and the heart was perfused retrogradely with a modified Krebs-Henseleit buffer (118.0 mM NaCl, 24.9 mM NaHCO_3_, 4.7 mM KCl, 1.75 mM CaCl_2_, 1.2 mM KH_2_PO_4_, 1.2 mM Mg_2_SO_4_, 11.0 mM glucose; 37 °C; pH = 7.4). The buffer was equilibrated with a mix of 5% CO_2_ and 95% O_2_. As soon as the heart was retrogradely perfused, the left atrium was cannulated and the heart was then placed in the heated chamber of the working-heart setup (Hugo Sachs Electronics, Germany) and perfused in Langendorff mode at a perfusion pressure of approximately 30 mmHg. Perfusion was then switched to antegrade, working-heart mode with inlet via the left atrium and ejection through the aorta. Left atrial filling pressure (preload) was varied between 10 and 25 mmHg. Afterload was set to 70 mmHg.

The left ventricular pressure was measured by a 0.9F optical pressure catheter (LS-PT9-10, FISO Technologies, Canada), placed in the left ventricle through the aorta. Two electrodes, positioned in a lead II configuration on the heart, recorded an electrocardiogram (ECG). A bipolar electrode was placed in direct contact with the right atrium to enable pacing of the heart at 8 Hz, using 1 ms current impulses provided from an isolated, external stimulator (DS3, Digitimer, Welwyn Garden City, UK). Hearts were discarded if they were unable to maintain a sinus rhythm above 300 bpm or a stroke volume at 10 mmHg filling pressure. In total, data from 8/12 control hearts and 9/12 HFpEF hearts were included in the study. 4 control hearts and 3 HFpEF hearts were excluded due to inability to maintain steady heart rhythm or leaks that was not possible to ligate adequately.

The following parameters were recorded continuously throughout the protocol: Temperature, ECG, left atrial pressure, left ventricular pressure, aorta pressure and aorta flow. Stroke volume was determined from the aorta flow curve and used for calculating cardiac output.[reference Usai] Peak left ventricular pressure, the peak slope of the ventricular relaxation (-dP/dt) and the ventricular diastolic time constant, τ, was calculated offline and blinded to group (HFpEF or control) in LabChart (Version 8.1.30, ADinstruments, Oxford, United Kingdom).

### Cardiomyocyte isolation

Hearts were explanted from mice rapidly following cervical dislocation and placed in ice-cold EDTA buffer (125 mM NaCl, 5 mM KCl, 0.5 mM NaH_2_PO_4_, 10 mM HEPES, 10 mM BDM, 10 mM taurine, 5 mM EDTA, 10 mM glucose; 37 °C; pH = 7.4). The explanted heart was carefully trimmed, the aorta cannulated and attached to a retrograde perfusion system with a peristaltic pump (KDS-100, KD Scientific). First the heart was perfused at 0.5 ml/min with 37 °C EDTA buffer for 6 minutes. Perfusion was then switched to a Perfusion buffer (130 mM NaCl, 5 mM KCl, 0.5 mM NaH_2_PO_4_, 10 mM HEPES, 10 mM BDM, 10 mM taurine, 1 mM MgCl_2_, 10 mM glucose; 37 °C; pH = 7.4) at 1.5 ml/min for 2 minutes. Finally, the heart was perfused with an enzyme solution (Perfusion Buffer containing 0.5 mg/mL collagenase II, 0.5 mg/mL collagenase IV and 0.05 mg/mL protease XIV) for 5-10 minutes until digestion is achieved. The left ventricle was cut off and tissue mechanically dissociated in Perfusion buffer before filtering the tissue solution through a fine mesh (300 μm filter) to isolate single cardiomyocytes. Cells were allowed to settle by gravity and then resuspended in Ca^2+^-free Tyrode’s solution (136 mM NaCl, 4 mM KCl, 5 mM HEPES, 5 mM MES, 0.8 mM MgCl_2_, 10 mM glucose; 37 °C; pH = 7.4). Calcium was introduced in a stepwise manner to the cell suspension over 30 minutes to a final concentration of 1.8 mM and stored at room temperature until use. In total, data from 4/9 control hearts and 8/12 HFpEF hearts were included in the study. In cases where cardiomyocyte isolation was unsuccessful, the yield of viable single cells was insufficient for experimental measurements, and these hearts were excluded from analyses.

### Cell morphometry

An aliquot of single cells in zero Ca^2+^ Tyrode were used for imaging from a subset of isolations (3 HFpEF and 3 control hearts). Images of cardiomyocytes were taken on a light microscope (Leica DMLB) using a 20X objective. Cardiomyocyte length, width and cross-sectional area were determined in a blinded-to-origin manner (Image Pro Premier, version 9).

### Calcium handling studies

The ratiometric Ca^2+^-sensitive dye Fura-2 AM (Invitrogen, Thermo Fisher) was used to monitor changes in [Ca^2+^]_i_. This ratiometric approach controls for variability in dye loading, photobleaching, and cell thickness, enabling reliable quantification of dynamic [Ca^2+^]_i_. To load myocytes, 2 μL of 1 μg/μL Fura-2 AM in DMSO was added to 1 mL cell suspension and protected from light. After 15 minutes, cells were resuspended in 1.8 mM Ca^2+^ Tyrode without Fura-2. Cells were left for a further 15 minutes to allow time for intracellular de-esterification of Fura-2 prior to analysis.

A single drop of cell solution was added to a laminin-coated glass coverslip inside a cell chamber (Warner Instruments) fitted with gravity-flow superfusion system with vacuum suction. Superfusate temperature was maintained using with a mTC3 inline heater (Cell MicroControls). Flow rate, adjusted by altering reservoir height, was set to 2.5 ml/min. The chamber was mounted on the stage of an inverted Olympus IX73 fluorescence microscope and the cells visualized with a ×40 air objective lens. A MyoPacer Field stimulator was used to apply 12V 3 ms bipolar pulse from platinum electrodes. Cells were superfused with 1.8 mM Ca^2+^ Tyrode at 37 °C. Cells with visually identifiable ultrastructural defects including membrane blebbing, lack of well-defined striations, cytoplasmic vesicles, automaticity or incompletely isolated cells were excluded from analyses. Calcium transients (CaTs) were recorded at stimulation frequencies between 0.5 and 4 Hz. A dual LED excitation system was used to excite cells alternately at 340 and 385 nm (250 Hz). Fura-2 emission >510nm detected via a photomultiplier tube was recorded in IonWizard (version 7.8.5.195, IonOptix). In a subset of cells, 50 nM isoprenaline was added to the superfusate.

Data were analysed blinded to group (HFpEF or control) offline. Diastolic Ca^2+^_i_ was defined asthe baseline fluorescence ratio of the two excitation wavelengths (F_340_/F_385_ or R_0_). Transient amplitude was defined as the peak Fura-2 fluorescence ratio (R) relative to R_0_ immediately preceding stimulation (R/R_0_). Time to peak was defined as the time interval from stimulation to the point at which the fluorescence ratio reached 90% of its maximum amplitude. Decay time was defined as the time from peak fluorescence to the point at which the fluorescence ratio had declined to 10% of its peak amplitude (i.e., 90% decay). Tau (τ), the time constant of Ca^2+^ transient decay, was calculated in IonWizard by fitting a single-exponential function to the declining phase of the transient from peak to baseline. τ corresponds to the time required for the fluorescence signal to fall to 36.8% (1/e) of its initial value at the start of decay.

### Statistics

All statistical analyses were performed using GraphPad Prism (version 10.5.0; GraphPad Software). Normality of data distributions was assessed using the Shapiro–Wilk test. For normally distributed data, differences between two groups were analysed using unpaired two-tailed Student’s *t*-tests; for non-normally distributed data, Mann–Whitney tests were applied. Multiple comparisons were corrected using the Holm–Šídák method where indicated. Repeated measures over time were analysed using mixed-effects models. Two-way ANOVA was used only for normally distributed data to evaluate the effects of group and preload in isolated heart experiments. For τ–frequency relationships, data were fitted with a one-phase decay model and compared using the sum-of-squares *F*-test. Data are reported as mean ± 95% confidence interval (CI) or median ± 95% CI as appropriate, and *p* < 0.05 was considered statistically significant.

## Acknowledgements

We thank Karin Larsen and Katrine Hegelund Rasmussen for their expert assistance with experiments and animal care.

